# Relative Hemodynamic Timing in Human White Matter

**DOI:** 10.64898/2026.05.29.728865

**Authors:** Muwei Li, Zhaohua Ding, John C Gore

**Author notes:** Corresponding author. Vanderbilt University Institute of Imaging Science, 1161 21st Ave. S, Medical Center North, AA-1105, Nashville, TN 37232-2310, USA.

## Abstract

Hemodynamic lag in white matter (WM) remains poorly understood despite its relevance to neurovascular health. We developed Local Propagation Mapping to quantify the spatiotemporal architecture of relative hemodynamic timing within WM. In young adults, WM lag architecture was reliable, aligned with venous anatomy, and reconfigured during working-memory task engagement; the magnitude of macroscopic lag modulation was associated with cognitive performance. Across aging, resting-state WM lag showed tract-specific bidirectional shifts and increased spatial fragmentation, with partial overlap with task-sensitive systems observed in young adults. Static baseline markers did not significantly mediate age-related cognitive differences, suggesting that aging effects may not operate through a simple resting-state pathway. Together, these findings establish relative WM lag as an informative dimension of neurovascular regulation and motivate a baseline-load hypothesis for future studies of cognitive aging.

## INTRODUCTION

Functional magnetic resonance imaging (fMRI) serves as the cornerstone for mapping human brain connectivity by capturing blood oxygen level-dependent (BOLD) signals. Because this intrinsic contrast mechanism relies on vascular hemodynamics rather than direct neural firing, inherent variations in blood transit time introduce critical temporal delays across distinct anatomical regions^1–4^. Previous investigations into this hemodynamic lag phenomenon have predominantly concentrated on the gray matter (GM) by tracking spontaneous propagation waves or by profiling systemic circulatory oscillations^5–7^. However, the spatiotemporal architecture of hemodynamic propagation within white matter (WM) remains largely uncharted. The lower baseline cerebral blood flow and distinct microvascular topology of WM tissue make accurate delay estimation particularly challenging. Consequently, the fundamental topographical organization of blood flow propagation along major anatomical tracts constitutes a major blind spot in current neurovascular research, severely limiting our understanding of how deep brain hemodynamics support global cognitive functions^6,8^.

Although arterial spin labeling (ASL) and hypercapnic challenges such as carbon dioxide inhalation are widely accepted as gold standards for evaluating cerebrovascular reactivity, they face insurmountable obstacles in WM research^9,10^. ASL measurements suffer from an inherently low signal-to-noise ratio in deep WM owing to prolonged transit times and reduced baseline perfusion^11^. Furthermore, hypercapnic experiments induce profound global vasodilation that inherently masks the subtle local neurovascular adjustments occurring during natural cognitive engagement^12,13^. The severe physiological stress associated with these gas challenges also makes them entirely incompatible with concurrent complex behavioral assessments. On the analytical front, traditional resting state delay estimation predominantly employs either global signal cross-correlation or GM reference seeding. Algorithms utilizing the global signal are heavily corrupted by systemic low-frequency fluctuations^2,14^. Because the contribution of these systemic components varies drastically across different tissue voxels, global lag estimations are often artificially biased by the local amplitude of the global component rather than reflecting true hemodynamic propagation^6,7,15^. Conversely, the alternative approach of using cortical GM or large venous structures as reference seeds introduces equally severe vulnerabilities^8,16^. This seed-based cross-correlation method inherently assumes a strong temporal synchrony between tissues. When applied to deep WM regions that lack robust functional connectivity with the remote GM reference, the mathematical correlation becomes dominated by stochastic noise, yielding entirely spurious lag estimations. The field, therefore, urgently requires a noninvasive and contrast-free methodology capable of precisely mapping the endogenous local propagation of hemodynamics within WM without imposing any physiological stress.

Beyond these methodological limitations lies a profound theoretical gap regarding how WM hemodynamics respond to transient cognitive demands versus chronic aging. During demanding mental operations, the human brain must dynamically reallocate blood flow resources. Yet it remains entirely unknown whether the deep WM vasculature possesses such state-dependent hemodynamic flexibility and how this transient spatiotemporal reconfiguration might relate to global cognitive efficiency. Conversely, healthy aging is known to severely alter baseline cerebrovascular responses^17–21^. However, the exact relationship between these chronic physiological drifts and cognitive decline remains deeply ambiguous. It is currently unclear whether age-related hemodynamic alterations directly mediate cognitive deterioration or merely represent parallel physiological weathering. Resolving this ambiguity requires a comprehensive framework to disentangle the acute functional compensations driven by cognitive tasks from the chronic physiological footprint imposed by aging.

To address these fundamental challenges, this study utilizes a large-scale dataset to introduce a novel Local Propagation Mapping (LPM) technique. By restricting cross-correlation analyses to immediate anatomical neighborhoods and solving a comprehensive Poisson equation, this methodology completely bypasses the spurious spatial correlations inherent to global reference approaches. We first demonstrate the physiological validity of this approach by establishing the superior reliability and the exceptional alignment with the macroscopic venous architecture of the brain. After characterizing the baseline hemodynamic gradients across seventy major WM tracts, we show that cognitive engagement expands the global temporal distribution and produces tract-specific reconfigurations in both macroscopic delay and spatial fragmentation. We then demonstrate that the magnitude of task-induced lag reconfiguration is associated with performance across multiple cognitive domains, suggesting that efficient cognition depends on economical and anatomically targeted hemodynamic tuning rather than large nonspecific redistribution^16,22^. Finally, we disentangle the effects of aging from those of task execution by showing that chronological age induces a localized chronic drift of the vascular baseline toward a configuration that resembles an active task state^17,18^. Our mediation analysis motivates a baseline-load model: static age-related biomarkers do not independently mediate cognitive decline, but their convergence with task-sensitive systems suggests that aging may narrow the efficiency margin available for task-evoked compensation. Ultimately, this work establishes a critical physiological baseline for WM hemodynamics and provides a unifying framework to understand neurovascular cognitive efficiency across the human lifespan.

## RESULTS

### Topographical Mapping, Reliability, and Physiological Validation of LPM

To address the limitations of traditional delay estimation, which predominantly employs either global signal cross-correlation or GM reference seeding and is often artificially biased by the local amplitude of noise, we developed the Local Propagation Mapping (LPM) approach. Briefly, LPM restricts cross-correlation analyses to immediate anatomical neighborhoods and solves a comprehensive Poisson equation. This methodology achieves sub-TR precision while completely bypassing the spurious spatial correlations inherent to global reference approaches.

As shown in Figure 1a-c, we compared group-averaged whole-brain hemodynamic lag maps derived from resting-state fMRI using our novel LPM method against two distinct analytical approaches: Global Signal Regression (GSR) lag mapping based on temporal correlation with the global mean signal, and Global Cross-Correlation (GCC) lag analysis, which estimates relative delays through exhaustive pair-wise correlations across all brain voxels. In these topographical maps, warm colors indicate delayed BOLD responses, whereas cool colors denote advanced responses. Crucially, we sought to biologically validate these estimations against an independent physiological reference, a macroscopic venous partial volume atlas where brighter colors reflect higher macroscopic venous density (Figure 1d). We examined the spatial association between estimated hemodynamic lags and underlying venous architecture. Voxels were parcellated into spatial bins based on their venous volume fractions. As shown in Figure 1g-i, scatter plots depicting the linear relationship between regional venous volume and the corresponding bin-averaged lag values revealed that LPM tightly aligns with the vascular anatomy (R^2^ = 0.948), vastly outperforming both GSR (R^2^ = 0.418) and GCC (R^2^ = 0.652). This physiological alignment was further confirmed via a subject-level contrast of estimated lags between regions of extremely high (top 25%) and low (bottom 25%) venous density. As shown in Figure 1j and k, across individual subject trajectories (N=687), LPM yielded the highest separation sensitivity, quantified as the effect size (Cohen’s d) of the high-versus-low venous volume lag contrast, highlighting the physiological specificity of the respective mapping approaches.

**Figure 1.**
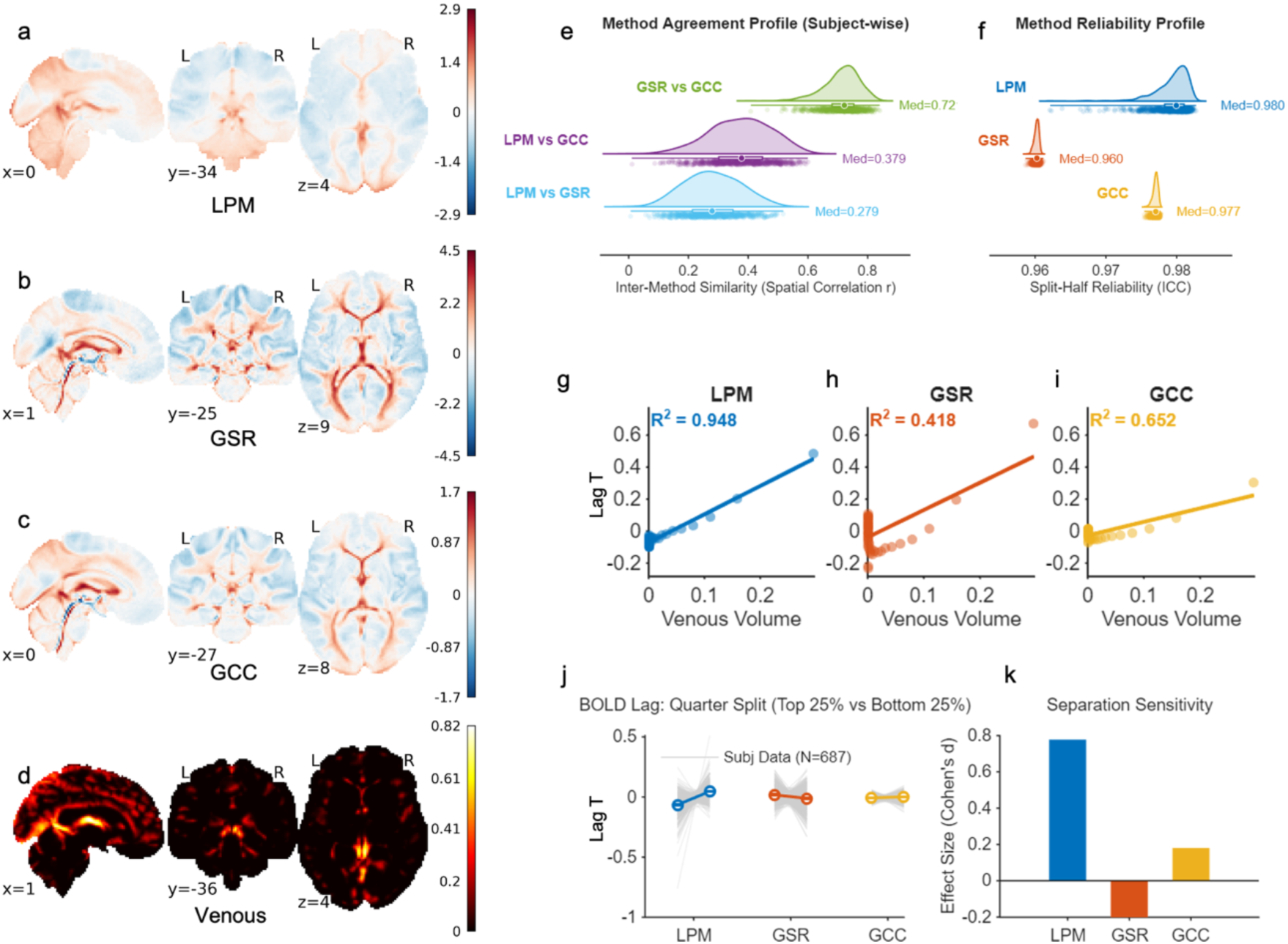
Topographical mapping, reliability, and physiological validation of hemodynamic lag estimation methods. (a-c) Group-averaged whole-brain hemodynamic lag maps derived from resting-state fMRI using three distinct analytical approaches: (a) our novel Local Propagation Mapping (LPM) method, (b) Global Signal Regression (GSR) lag mapping based on temporal correlation with the global mean signal, and (c) Global Cross-Correlation (GCC) lag mapping analysis. Warm colors indicate delayed BOLD responses, whereas cool colors denote advanced responses. (d) A macroscopic venous partial volume atlas is used as an independent physiological reference. Brighter colors reflect higher macroscopic venous density. (e) Inter-method spatial agreement. Raincloud plots show the distribution of subject-wise spatial correlations (Pearson’s r) between the lag maps generated by the three methods. (f) Split-half reliability of each method, quantified by the spatial Intraclass Correlation Coefficient (ICC). Distributions reflect robust intra-method consistency across iterative half-splits. (g-i) Spatial association between estimated hemodynamic lags and underlying venous architecture. Voxels were parcellated into spatial bins based on their venous volume fractions. Scatter plots depict the linear relationship between regional venous volume and the corresponding bin-averaged lag values for (g) LPM, (h) GSR, and (i) GCC, along with the coefficient of determination R^2^. (j) Subject-level contrast of estimated delays between regions of extremely high (top 25%) and low (bottom 25%) venous density. Gray lines trace individual subject trajectories (N = 687), with colored markers indicating group means. (k) Separation sensitivity, quantified as the effect size (Cohen’s d) of the high-versus-low venous volume lag contrast, highlighting the physiological specificity of the respective mapping approaches.

We evaluated the split-half reliability of each method, quantified by the spatial Intraclass Correlation Coefficient (ICC). As shown in Figure 1f, LPM demonstrated exceptional stability, with distributions reflecting robust intra-method consistency across iterative half-splits. Furthermore, raincloud plots showing the distribution of subject-wise spatial correlations (Pearson’s r) mapped the inter-method spatial agreement between the lag maps generated by the three methods (Figure 1e).

### Spatial Profiling of Hemodynamic Propagation in WM Tracts

To characterize the spatiotemporal architecture of hemodynamic propagation within the WM, we projected the LPM-derived delay maps onto a standard anatomical atlas. Using deterministic fiber tractography, we extracted hemodynamic Lag signals (Lag T) at 100 equidistant nodes along the trajectories of 70 major WM bundles. This along-tract analysis effectively captures the macroscopic lag gradients within specific neural pathways.

Our analysis revealed that WM hemodynamics are not uniform but follow highly organized spatial gradients that align with the underlying axonal architecture. For instance, in the left corticospinal tract (CST L), we used three-dimensional topographical mapping and the along-tract profile to show the clear progression of delays (Figure 2a). Earlier BOLD responses (negative Lag T) are concentrated in inferior/deep regions, while delays progressively increase as the tract extends toward superior cortical terminations. Expanding this analysis to all 70 major WM bundles confirmed the universality of these structured gradients (Figure 2b). Each tract exhibits a distinct and highly conserved spatial “fingerprint” of hemodynamic delays.

**Figure 2.**
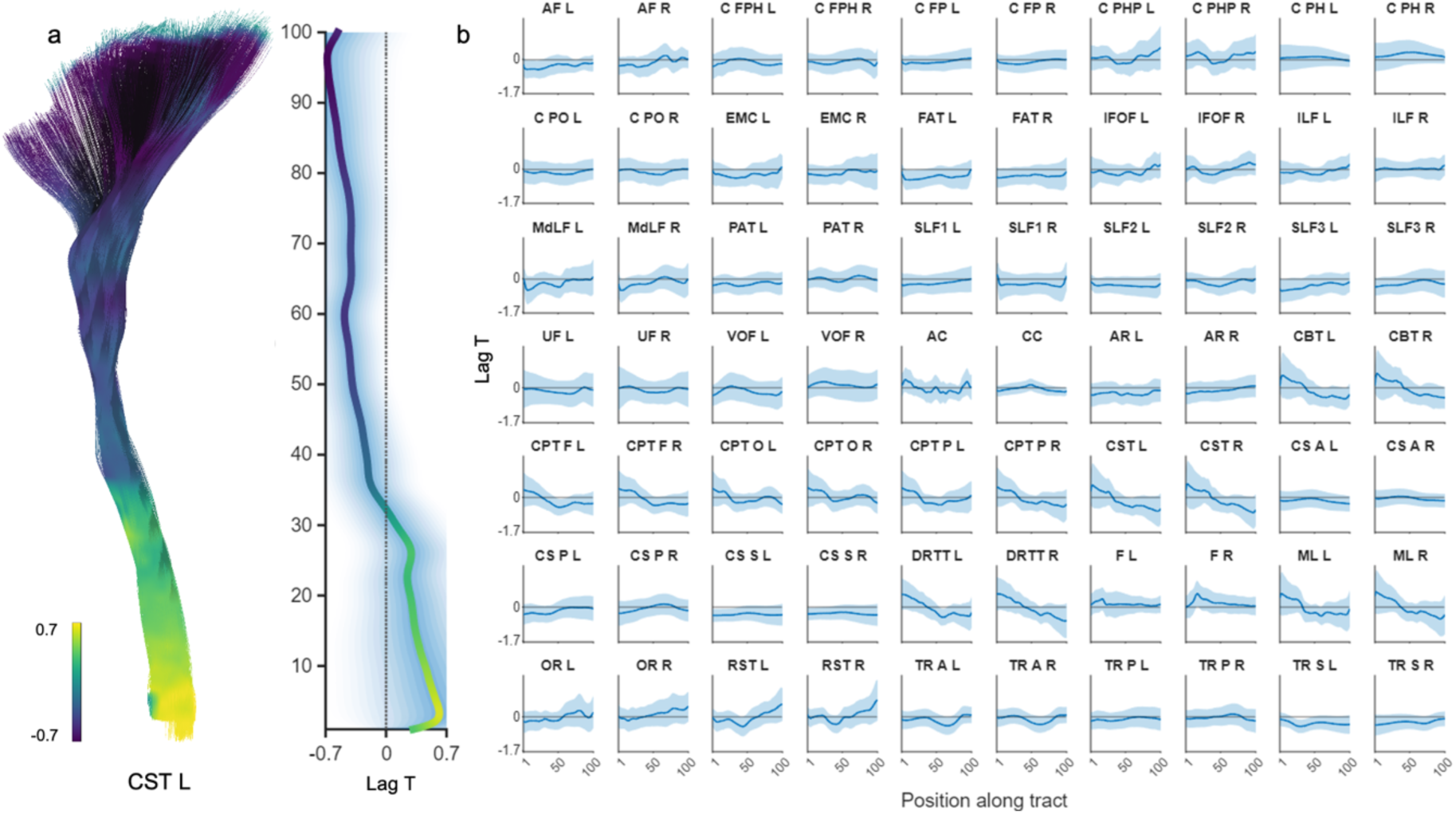
Spatial profiling of hemodynamic Lags along major white matter tracts. (a) Three-dimensional topographical mapping of hemodynamic lag (Lag T) along the left corticospinal tract (CST L). Streamlines are color-coded according to local delay values, with cooler colors indicating earlier BOLD responses (negative Lag T) and warmer colors denoting delayed responses (positive Lag T). The adjacent vertical line plot displays the corresponding along-tract hemodynamic profile. The trajectory traces the spatial evolution of Lag T across 100 equidistant nodes, progressing from inferior/deep regions (bottom) to superior/cortical terminations (top), with the shaded envelope representing inter-subject variability (s.d.). (b) Comprehensive along-tract hemodynamic profiles for 70 major white matter bundles. Each subpanel illustrates the distinct spatial gradient of delays specific to the anatomical trajectory of the tract (x-axis: standardized position along the tract from node 1 to 100; y-axis: Lag T). Solid blue lines denote the group-level mean, and shaded regions represent the standard deviation across the cohort.

### Task-Induced Modulations of Hemodynamic Lags

Having established the baseline spatial gradients of WM hemodynamics, we investigated whether these deep vascular networks possess state-dependent reconfigurability. We compared the hemodynamic architecture during the resting state to that during a cognitively demanding working memory task, quantifying these modulations from the global whole-brain scale down to specific anatomical tracts.

To assess global hemodynamic shifts, we first examined the voxel-wise whole-brain WM lag distributions (Figure 3a). A group-level statistical comparison of the Full Width at Half Maximum (FWHM) revealed a highly significant task-induced widening of this temporal gradient (Figure 3b, P < 0.001). This global expansion indicates that cognitive engagement broadens the dispersion of relative hemodynamic timing across the WM.

**Figure 3.**
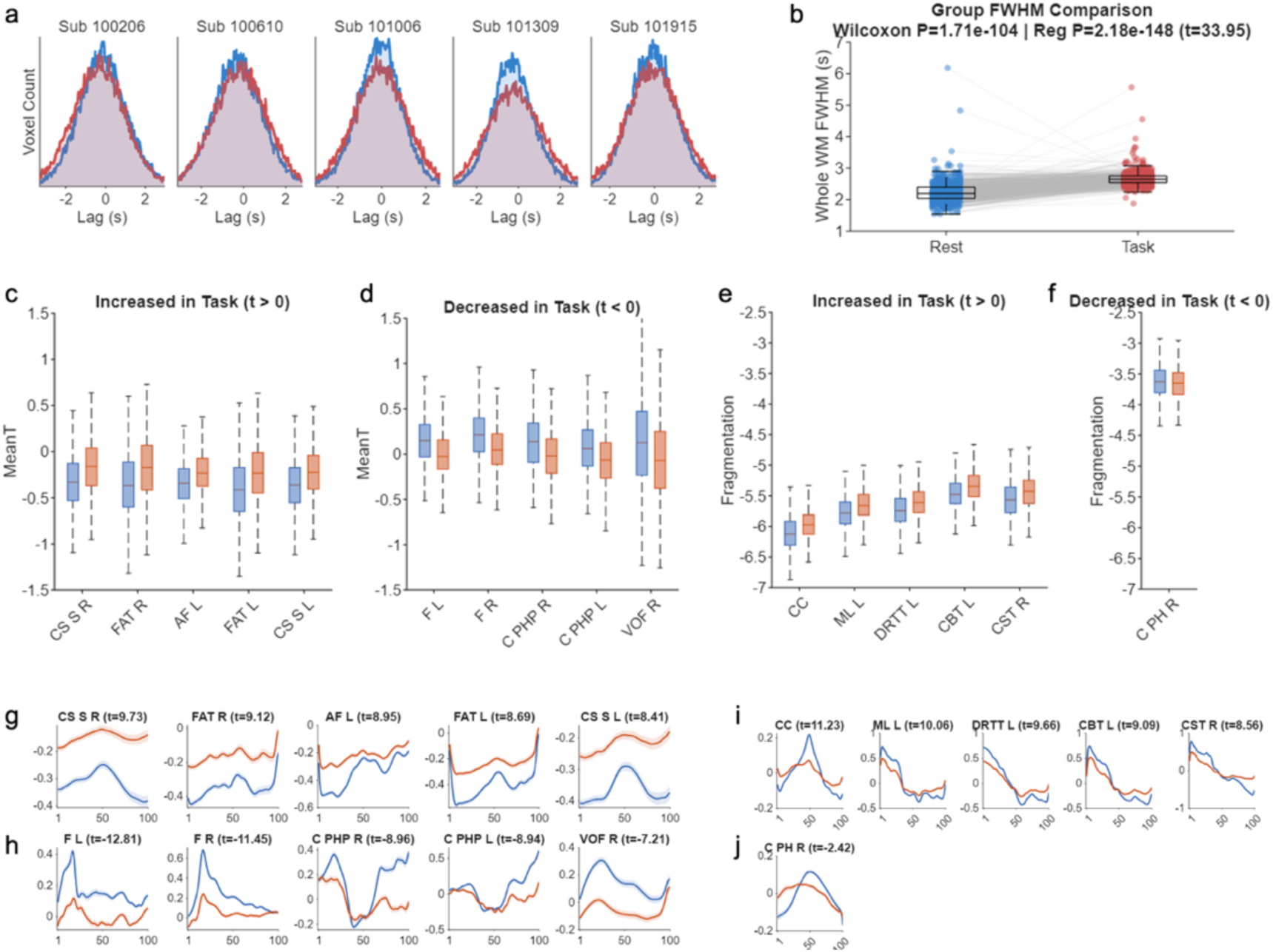
Task-induced modulations of hemodynamic lags in WM. (a) Whole-brain white matter lag distributions. Voxel-wise histograms of hemodynamic delay (Lag) across the entire white matter for five representative subjects. Distributions during the resting state are shown in blue, and those during the working memory task are shown in red. (b) Group-level FWHM comparison. Statistical comparison of the Full Width at Half Maximum (FWHM) of the whole white matter lag distribution between rest and task states across all subjects. The task state exhibits a significantly wider FWHM (P < 0.001), indicating a task-induced functional segregation and widening of the temporal hemodynamic gradient. Boxplots display the interquartile range and median; individual subject trajectories are connected by light grey lines. Significance was assessed using both a paired Wilcoxon signed-rank test and a linear regression model controlling for the difference in head motion. (c–f) Group-level comparison of hemodynamic properties between resting-state (blue boxes) and task-state (orange boxes). Boxplots illustrate the top 5 most significantly altered white matter tracts showing task-induced increases (left columns) and decreases (right columns) across two distinct metrics: (c, d) Macroscopic delay (Mean T), reflecting overall temporal shifts in blood flow, and (e, f) Spatial fragmentation (spectral slope β), capturing the microscopic spatial complexity of the propagation pattern. Note that due to stringent statistical thresholding (FDR-corrected P < 0.05, adjusting for state-specific head motion), only one tract exhibited a significant decrease in fragmentation during the task (f). (g–j) Detailed along-tract spatial profiles corresponding to the significant tracts identified above. Panels (g) and (h) trace the absolute Mean T trajectories across 100 equidistant nodes for the tracts in (c) and (d), revealing where along the anatomical bundle the task-induced acceleration or deceleration occurs. Solid lines denote the group means (blue: rest, orange: task), with shaded envelopes representing the standard error of the mean (s.e.m.). Panels (i) and (j) depict the spatially demeaned Lag T signals for the tracts in (e) and (f).

Zooming into the along-tract level, we profiled the precise spatial alterations across the 70 major WM bundles using two complementary metrics: macroscopic delay (Mean T), reflecting the overall relative temporal position of each tract within the lag field, and spatial fragmentation (spectral slope β), capturing the microscopic complexity of the propagation waveform. After strictly controlling for state-specific head motion and applying False Discovery Rate (FDR) correction, we observed highly structured, tract-specific neurovascular reconfigurations. For macroscopic delays, the task induced significant bidirectional shifts: certain tracts shifted toward later relative timing (Figure 3c), while others shifted toward earlier relative timing (Figure 3d). The detailed spatial gradients of these top tracts reveal precisely where along the anatomical bundle these relative task-rest shifts occur (Figure 3g, h). In contrast, the microscopic spatial fragmentation exhibited a strongly unidirectional global trend. We observed widespread task-induced increases in spatial fragmentation across the WM architecture (Figure 3e, i), indicating that cognitive operations require a more complex, high-frequency spatial regulation of local hemodynamics. Notably, only a single tract exhibited a significant decrease in fragmentation during the task (Figure 3f, j). An exhaustive profiling of these task-induced modulations across all significant WM tracts is provided in Supplementary Figure S1.

### Task-Induced Hemodynamic Reconfiguration is Associated with Cognitive Performance

We next tested whether the magnitude of this task-induced vascular reconfiguration is behaviorally relevant. To this end, we calculated the absolute task-rest difference in hemodynamic lag (ΔLag = |Lagtask - Lagrest|) for each tract and evaluated its association with individual performance across multiple cognitive domains.

To isolate the unique contribution of task-induced lag reconfiguration, we employed linear regression models that rigorously partialed out the confounding effects of chronological age, biological sex, baseline resting-state lag, and the difference in head motion between the two states. The resulting cross-domain analysis (Figure 4) revealed widespread, significant associations (P_FDR_ < 0.05) between tract-specific ΔLag and diverse cognitive abilities, including fluid intelligence, working memory accuracy, processing speed, and cognitive flexibility (card sorting). The directionality of these relationships, indicated by the blue (negative) and orange (positive) T-statistics in the heatmap, highlights a complex neural-efficiency mechanism. In many domains, larger task-rest perturbations were associated with lower cognitive performance, suggesting that better performance may depend on more economical and targeted hemodynamic tuning rather than larger vascular redistribution. To illustrate the robustness of these associations at the individual subject level, partial regression scatter plots isolating the unique variance for the most significant tracts across each cognitive domain are detailed in Supplementary Figure S2.

**Figure 4.**
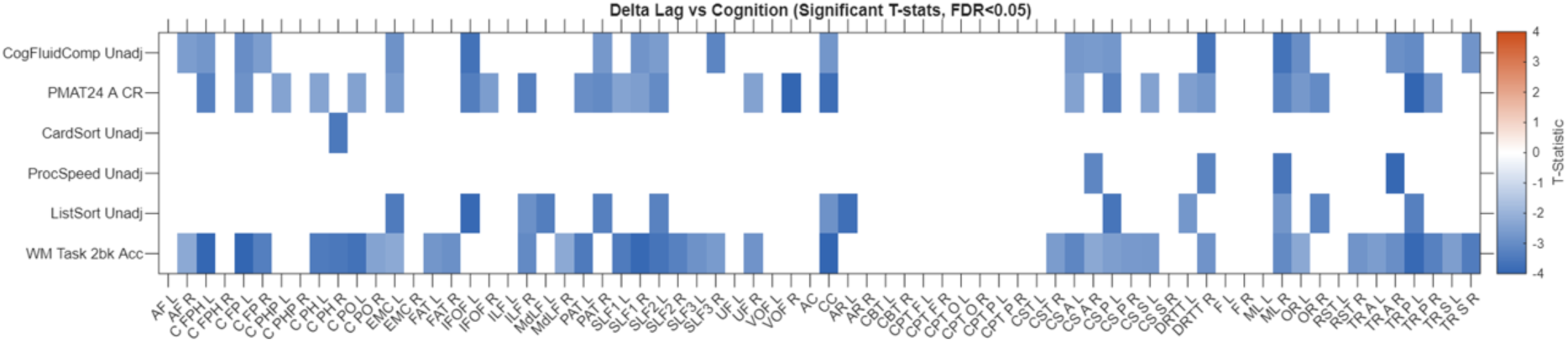
Relationship between task-induced hemodynamic delay modulation and cognitive performance. The heatmap displays the T-statistics from linear regression models relating the absolute task-rest difference in hemodynamic lag (Delta Lag = |Lag_task_ - Lag_rest_|) to various cognitive and behavioral scores across white matter tracts. Each cell represents the association for a specific tract (x-axis) and cognitive domain (y-axis). All models controlled for age, sex, baseline lag (resting state), and the difference in head motion between task and resting states. White cells indicate non-significant associations after False Discovery Rate (FDR) correction (P_FDR_ > 0.05). The color bar represents the T-statistic, where blue indicates a negative correlation (a larger Delta Lag associated with lower cognitive performance), and orange indicates a positive correlation.

While the spatial fragmentation (β) of the hemodynamic waveform was significantly altered by the task at the group level, individual differences in the magnitude of this shift (Δβ) did not exhibit significant associations with cognitive performance across the tested domains.

### Changes in Hemodynamic Delay with Aging

Having demonstrated acute state-dependent reconfiguration of WM hemodynamics, we further evaluated the chronic drift of this baseline architecture during healthy aging. Cross-sectional analysis revealed that the macroscopic hemodynamic delay (Mean T) undergoes significant and anatomically specific spatial reorganization as a function of chronological age (Figure 5). This aging effect exhibited a highly structured bidirectional pattern. As detailed in Table 1, 22 distinct WM tracts demonstrated significant age-related modulations (FDR-corrected P < 0.05). Specifically, certain pathways, such as the left frontal aslant tract (FAT L, t = 4.59) and the left anterior corticostriatal tract (CS A L, t = 3.96), showed a robust positive age effect, characterized by a progressive shift toward later relative lag with advancing age (Figure 5a). Conversely, pathways including the bilateral optic radiations (OR L, t = -4.74; OR R, t = -4.09) exhibited strong negative age effects, characterized by progressive shifts toward earlier relative lag (Figure 5b). This bidirectional drift indicates that aging does not uniformly slow down WM hemodynamics, but rather redistributes relative baseline timing across different anatomical networks.

**Figure 5.**
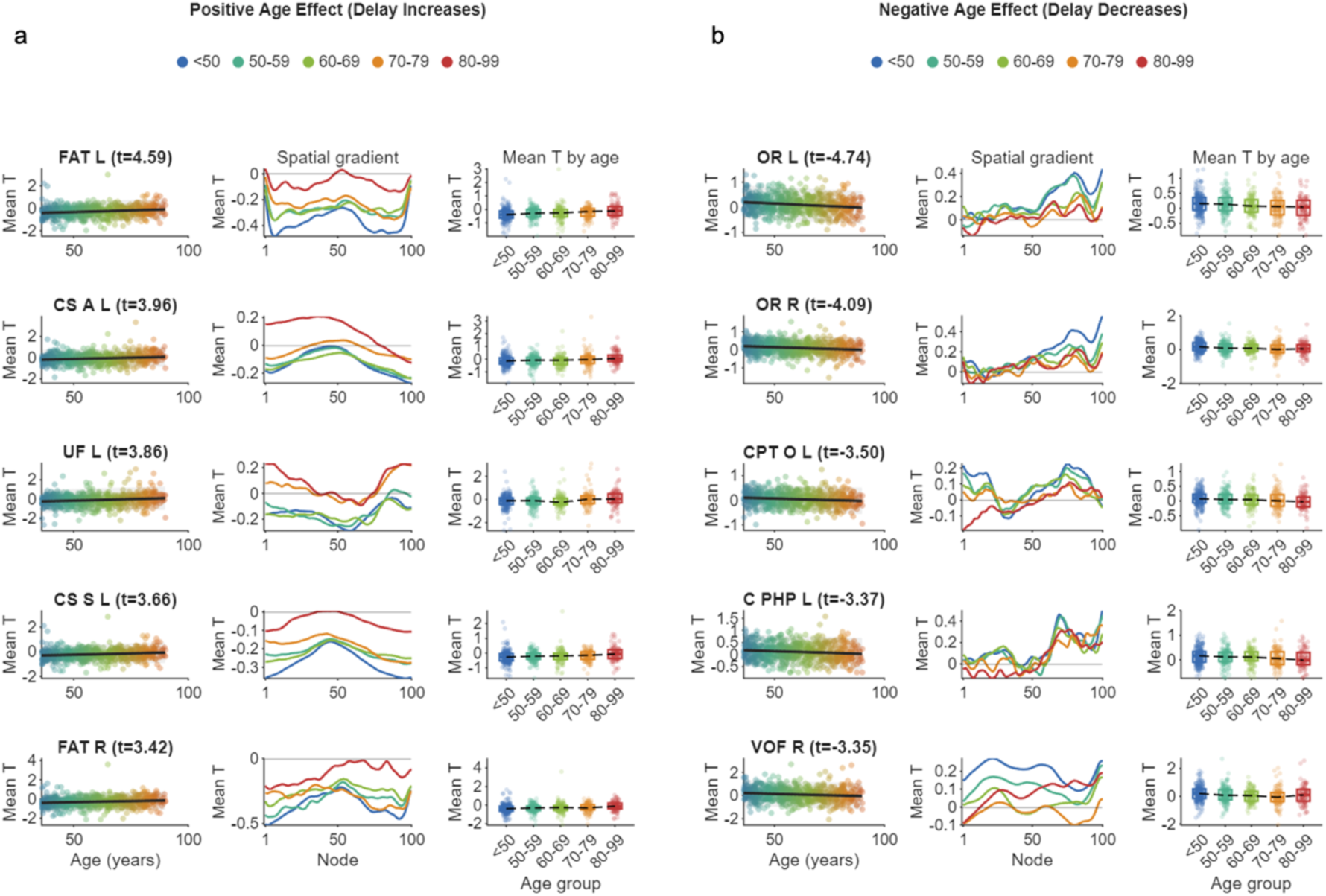
Aging-associated alterations in the hemodynamic Lags. Age-related modulations of macroscopic hemodynamic Lags (Mean T) across major white matter tracts. Significant age associations (FDR-corrected P < 0.05, adjusting for sex and head motion covariates) are stratified into (a) Positive age effects (tracts showing progressive delay increases with age, t > 0) and (b) Negative age effects (tracts showing progressive delay decreases with age, t < 0). For each significant tract, three distinct visualizations are provided (from left to right): (1) A scatter plot displaying the continuous linear relationship between chronological age (years) and the tract-averaged Mean T. The solid black line represents the linear regression fit. (2) Along-tract spatial gradients (from node 1 to 100) averaged across five decades of life (color-coded from <50 years in blue to 80-99 years in red). These profiles reveal how the spatial waveform of the hemodynamic propagation progressively shifts across the human lifespan. (3) Categorical distributions of the tract-averaged Mean T across the same five age groups, presented as combined box-and-scatter plots. The dashed black line connects the group medians, clearly illustrating the discrete trajectory of the aging effect. The respective t-statistic representing the independent main effect of age is indicated adjacent to each anatomical tract abbreviation.

**Table 1.** Significant age effects on mean Lags (FDR < 0.05)

| Tract Name | t-statistic | p-uncorrected | p-FDR |
| --- | --- | --- | --- |
| FAT_L | 4.59 | 5.34e-06 | 1.87e-04 |
| CS_A_L | 3.96 | 8.29e-05 | 0.0015 |
| UF_L | 3.86 | 1.26e-04 | 0.0018 |
| CS_S_L | 3.66 | 2.71e-04 | 0.0032 |
| FAT_R | 3.42 | 6.54e-04 | 0.0057 |
| C_PO_R | 3.19 | 0.0015 | 0.0094 |
| TR_S_L | 3.16 | 0.0016 | 0.0096 |
| TR_A_L | 2.98 | 0.0030 | 0.0162 |
| EMC_L | 2.92 | 0.0036 | 0.0180 |
| CS_S_R | 2.87 | 0.0043 | 0.0195 |
| CS_A_R | 2.85 | 0.0045 | 0.0195 |
| C_PO_L | 2.71 | 0.0070 | 0.0281 |
| AF_L | 2.69 | 0.0072 | 0.0281 |
| UF_R | 2.64 | 0.0085 | 0.0314 |
| TR_A_R | 2.48 | 0.0134 | 0.0427 |
| ILF_L | -2.52 | 0.0120 | 0.0400 |
| VOF_L | -2.55 | 0.0109 | 0.0383 |
| VOF_R | -3.35 | 8.62e-04 | 0.0060 |
| C_PHP_L | -3.37 | 8.08e-04 | 0.0060 |
| CPT_O_L | -3.50 | 4.94e-04 | 0.0049 |
| OR_R | -4.09 | 4.84e-05 | 0.0011 |
| OR_L | -4.74 | 2.62e-06 | 1.83e-04 |

Note: Results are sorted by the value of the t-statistic. All listed tracts survived False Discovery Rate (FDR) correction at $p < 0.05$ . L: Left hemisphere; R: Right hemisphere.

Notably, despite these local baseline shifts within specific tracts, the overall dispersion of the whole-brain WM hemodynamic gradient showed no evidence of age-related change. An evaluation of the FWHM of the global delay distribution revealed no significant correlation with age (P = 0.936), and the normalized frequency distributions of different age cohorts were highly similar (Supplementary Figure S3). These findings do not establish an active global balancing mechanism; rather, they indicate that aging-related effects in this dataset are primarily local and tract-specific rather than reflecting a uniform expansion or contraction of the global lag distribution.

### Progressive Fragmentation of Microscopic Spatial Complexity in Aging

Beyond macroscopic spatial shifts, we quantified the age-related alterations in the microscopic spatial complexity of the propagating waveforms using the spatial spectral slope. As chronological age advanced, the hemodynamic spatial gradients within specific tracts became increasingly fragmented (Figure 6). Statistical modeling identified significant age-related fragmentation, indicated by progressively shallower spectral slopes, in six key tracts (FDR-corrected P < 0.05). As summarized in Table 2, this effect was most prominent in the bilateral frontal parahippocampal cingulum (C FPH R, t = 3.71; C FPH L, t = 3.36) and the corpus callosum (CC, t = 3.07). Visualizing the spatially demeaned gradients across distinct age decades confirmed that older cohorts (80-99 years) exhibit substantially higher-frequency spatial fluctuations compared to younger individuals (<50 years) (Figure 6, middle panels).

**Figure 6.**
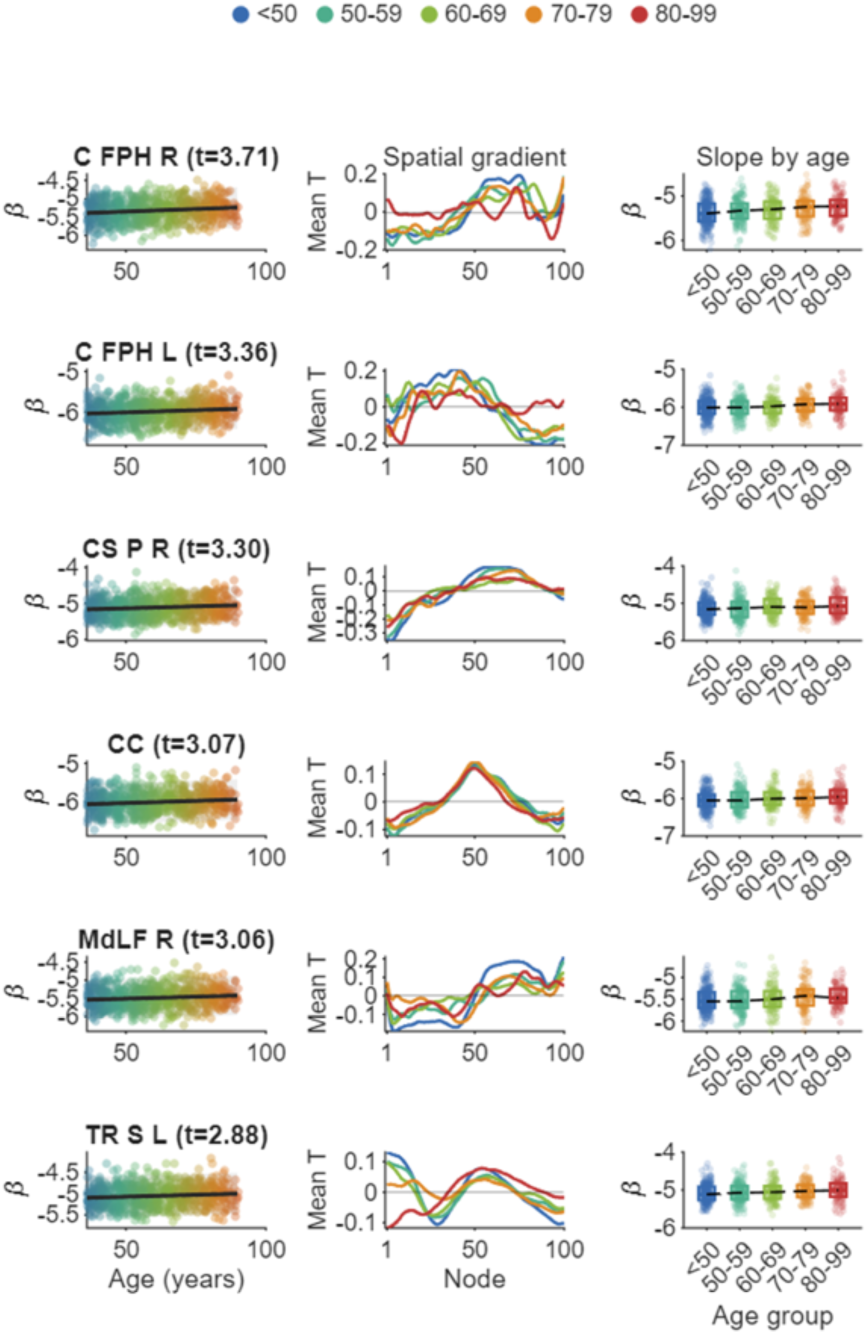
Aging-associated alterations in the spatial fragmentation of hemodynamic Lags. Age-related modulations of microscopic spatial complexity across major white matter tracts. Significant age associations (FDR-corrected P < 0.05, adjusting for sex and head motion covariates) are shown for tracts exhibiting progressive age-related changes in β. For each significant tract, three distinct visualizations illustrate the trajectory of the aging effect (from left to right): (1) A scatter plot depicting the continuous linear relationship between chronological age (years) and the tract-level spectral slope β. The solid black line indicates the linear regression fit. (2) Spatially demeaned along-tract lag profiles (from node 1 to 100) averaged within distinct aging decades (color-coded from <50 years in blue to 80-99 years in red). These spatial gradients visually demonstrate the age-dependent alterations in the high-frequency spatial fluctuations (fragmentation) of the propagating hemodynamic waves. (3) Categorical distributions of the spectral slope β across the five age groups, presented as combined box-and-scatter plots. The dashed black line connects the group medians, clearly illustrating the discrete stepwise progression of the aging effect. The respective t-statistic representing the independent main effect of age is indicated adjacent to each anatomical tract abbreviation.

**Table 2.** Statistical Results: Aging Effect on Lag Fragmentation (FDR < 0.05)

| Tract Name | t-statistic | p-uncorrected | p-FDR |
| --- | --- | --- | --- |
| C_FPH_R | 3.71 | 2.29e-04 | 0.0160 |
| C_FPH_L | 3.36 | 8.12e-04 | 0.0238 |
| CS_P_R | 3.30 | 0.0010 | 0.0238 |
| CC | 3.07 | 0.0022 | 0.0318 |
| MdLF_R | 3.06 | 0.0023 | 0.0318 |
| TR_S_L | 2.88 | 0.0042 | 0.0486 |

Note: Results are sorted by the value of the t-statistic. All listed tracts survived False Discovery Rate (FDR) correction at $p < 0.05$ .

Taken together with the task results, this chronic, age-induced increase in baseline spatial fragmentation partially resembles the acute lag reconfiguration observed during cognitive load. This convergence was most apparent for spatial fragmentation: five of the six age-sensitive tracts (C FPH R, C FPH L, CS P R, CC, and MdLF R) also showed task-induced increases in fragmentation. A similar but less complete pattern was observed for macroscopic Mean T, where 13 of the 22 age-sensitive tracts showed task-related shifts in the same direction, including overlapping positive shifts in AF L, EMC L, FAT L/R, CS S L/R, and TR S L, and overlapping negative shifts in C PHP L, CPT O L, VOF L/R, and OR L/R. This anatomical concordance suggests that aging may shift the resting cerebrovascular architecture toward a more task-like, higher-load configuration in selected WM systems. Because task-induced lag reconfiguration was behaviorally meaningful, such elevated baseline complexity may reduce the efficiency margin available for economical task-evoked regulation, although the present study does not directly measure age-related changes in task-evoked dynamic capacity.

### Dissecting the Bottleneck in the Mediation of Age-Related Cognitive Decline

To elucidate whether the profound age-related physiological and structural shifts directly drive cognitive deterioration, we performed a comprehensive mediation analysis. We evaluated the sequential pathways linking chronological age, neuroimaging biomarkers, and individual performance across three distinct cognitive domains (Flanker, Card Sort, and List Sort). We systematically compared three WM properties: macroscopic hemodynamic delay (Mean T), microscopic spatial complexity (Fragmentation), and structural myelin content (derived from T1w/T2w ratios). The ten anatomical tracts incorporated into this analysis were specifically selected based on their peak sensitivity to aging in the spatial fragmentation metric.

Our analysis revealed a striking dissociation within the mediation pathways (Figure 7). For Path A (Sensitivity to Aging, Age → Brain), structural myelin content and hemodynamic spatial fragmentation exhibited robust and highly significant change with advancing age, easily exceeding the nominal significance threshold (Z > 1.96) (A panels), confirming that the deep WM undergoes severe chronic physiological weathering.

**Figure 7.**
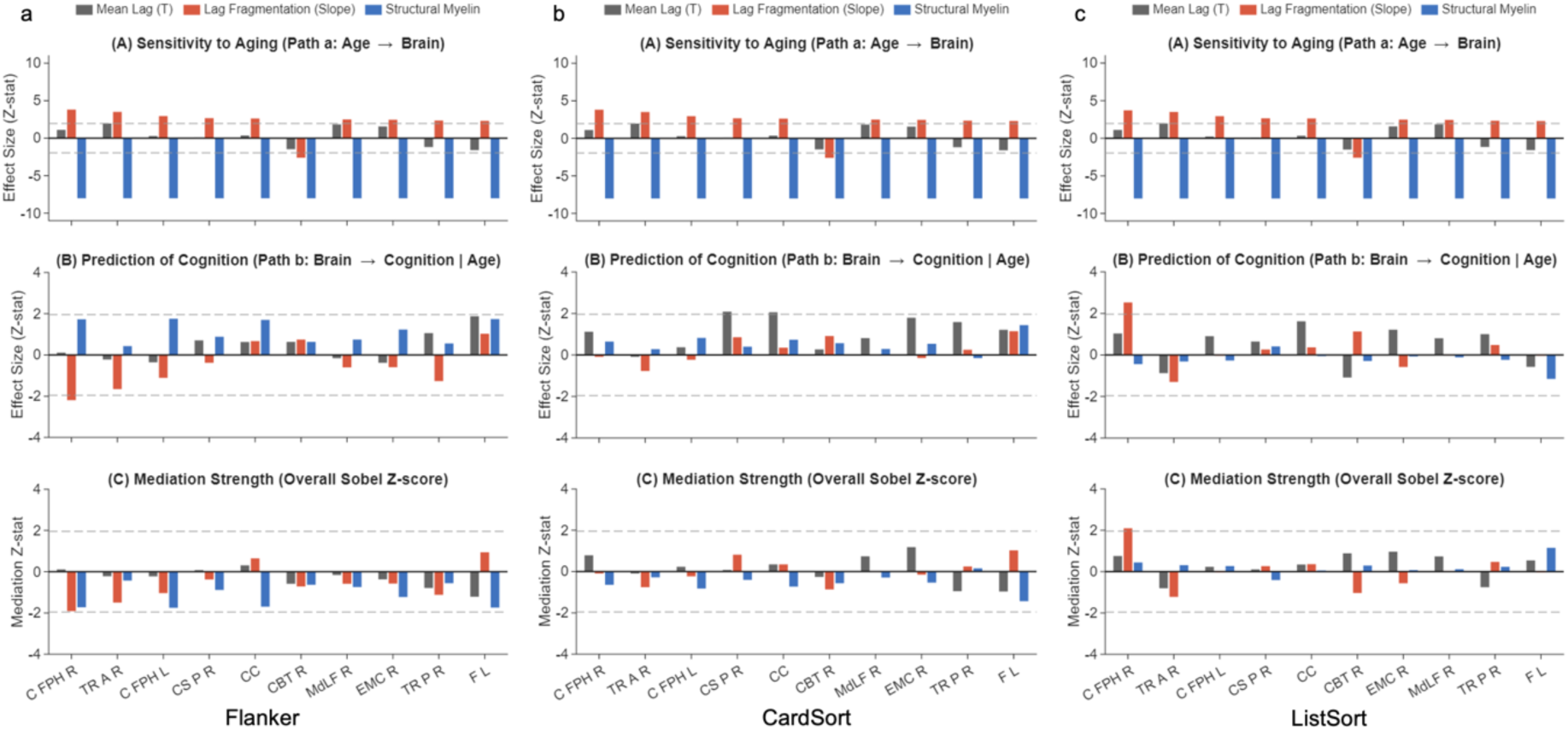
Dissecting the bottleneck in the mediation of age-related cognitive decline by white matter biomarkers. Comparative mediation analysis evaluating the sequential pathways linking chronological age, neuroimaging biomarkers, and performance across three distinct cognitive domains (columns: Flanker, CardSort, ListSort). Three white matter properties are compared: macroscopic hemodynamic delay (Mean T, gray), microscopic hemodynamic complexity (Lag Fragmentation, red), and structural myelin content (blue). The ten anatomical tracts displayed on the x-axis were specifically selected based on their peak sensitivity (top 10) to aging in the fragmentation metric. (A) Sensitivity to Aging (Path a: Age -> Brain). Bar heights represent the Z-statistic of the age effect. Both structural myelin and lag fragmentation exhibit robust, highly significant associations with age, easily exceeding the nominal significance threshold (dashed gray lines, Z = 1.96). (B) Prediction of Cognition (Path b: Brain -> Cognition | Age). The independent predictive value of each brain marker for cognitive performance, after statistically regressing out the variance of chronological age. Strikingly, despite the profound age-related physiological changes observed in (A), none of the markers reliably predict cognitive performance; almost all effect sizes collapse and fail to breach the significance threshold. (C) Overall Mediation Strength (Sobel Z-score). Due to the predictive bottleneck at Path b, the combined mediation effects fall short of significance across all markers and tasks.

However, a critical “predictive bottleneck” emerged at Path B (Prediction of Cognition, Brain → Cognition | Age). We assessed the independent predictive value of each brain marker for cognitive performance after statistically regressing out the shared variance of chronological age. Strikingly, despite the profound age-related physiological changes established in Path A, none of the baseline structural or hemodynamic markers reliably predicted cognitive performance (B panels). Almost all effect sizes collapsed, failing to breach the significance threshold across all three cognitive domains.

Consequently, the overall mediation strength (quantified by the Sobel Z-score) fell short of significance across all tested markers and cognitive tasks (C panels). These results indicate that the static, chronic physiological footprint of aging (baseline weathering) does not independently mediate cognitive decline. Integrating these findings with our task-state results (Figure 4), we propose a more cautious interpretation: baseline vascular degradation may affect cognition not through a simple static mediation pathway, but by shifting the system toward a high-load resting configuration that could constrain efficient task-evoked reconfiguration. This baseline-load mechanism remains a hypothesis to be tested directly in datasets with comparable task paradigms across the adult lifespan.

## DISCUSSION

In this study, we mapped the elusive spatiotemporal architecture of hemodynamic lag within the human WM to uncover its functional significance in cognitive efficiency and aging. By developing and physiologically validating the LPM approach, we overcame the limitations of traditional global-reference techniques to accurately trace relative hemodynamic timing along major anatomical pathways. We discovered that WM hemodynamics exhibit highly structured, tract-specific spatial gradients that are not merely passive physiological pipelines, but dynamically responsive networks. During cognitive engagement, these deep vascular pathways undergo profound spatiotemporal reconfigurations, and the magnitude of this task-induced lag reconfiguration is associated with individual performance across diverse cognitive domains. Notably, the predominance of negative associations suggests that better performance may be supported by more economical hemodynamic tuning rather than larger task-rest perturbations. Furthermore, we revealed that healthy aging imposes a chronic, bidirectional remodeling of this baseline architecture and progressively fragments its spatial complexity, partially resembling the acute physiological demands of high cognitive load. Crucially, our mediation analysis identified a predictive bottleneck: while aging severely alters both structural myelin and baseline hemodynamics, these static physiological footprints do not independently drive cognitive deterioration. Together, these findings suggest that relative WM hemodynamic timing is dynamically organized and behaviorally relevant, and they motivate a baseline-load hypothesis in which aging-related resting-state alterations may constrain task-evoked regulation.

### Reimagining WM: From Nuisance Regressors to Functional Networks

For decades, the neuroscientific community has operated under a predominantly “corticocentric” paradigm, where BOLD fluctuations in WM were systematically dismissed as instrumental artifacts or physiological noise^23–25^. This skepticism was rooted in the lower vascular density and metabolic demands of WM compared to the cortex. However, our findings, facilitated by the high-precision LPM, join a transformative body of evidence suggesting that WM is an active and organized component of the functional brain.

The physiological validity of our along-tract hemodynamic profiles is strongly supported by recent evidence demonstrating that WM-BOLD signals are not random but reflect underlying neural activity^21,26–32^. Specifically, intracranial electrophysiological recordings have confirmed that BOLD functional connectivity in WM correlates with local field potential synchronization^33^. This suggests that the spatiotemporal gradients we observed are the macroscopic manifestations of deep-seated neurovascular coupling, rather than simple partial-volume effects from the adjacent GM. Furthermore, our characterization of state-dependent reconfigurability in WM aligns with emerging theories of the “three-way” functional connectome^34^. The integration of WM as a functional “edge” rather than a passive “pipe” provides a missing third dimension to our understanding of brain networks. By demonstrating that task engagement triggers a structured reconfiguration of along-tract propagation, we provide empirical support for the notion that WM pathways actively modulate information transmission in response to cognitive demand^19,35^.

The chronic fragmentation of these gradients observed in aging echoes previous findings of disrupted WM functional organization in pathology, such as Alzheimer’s disease and schizophrenia^36,37^. Our study elevates this discussion by suggesting that the “weathering” of WM function is a progressive shift in the spatial spectral properties of hemodynamics. The aging-related fragmentation observed here also resonates with prior reports of disrupted WM functional organization in aging and neuropsychiatric conditions. This convergence motivates a more specific interpretation of lag fragmentation and baseline load, developed below.

### Distinct Roles of System-Level Lag Tuning and Local Spatial Fragmentation

Our findings suggest that WM hemodynamic lag is regulated at two related but distinct spatial scales. At the tract level, mean lag reflects the relative temporal position of a white matter pathway within the broader lag field. In contrast, the spectral slope captures how smooth or irregular the lag profile is along the tract. Both measures were sensitive to cognitive state and aging, indicating that WM hemodynamics are not static. However, task-induced changes in mean lag showed the clearest relationship with individual cognitive performance, whereas changes in spatial fragmentation did not show comparable behavioral associations.

This dissociation suggests that local spatial fragmentation and system-level lag tuning may reflect different aspects of vascular regulation. Increased fragmentation, expressed as a shallower spatial power spectrum, indicates that the lag profile becomes less smooth and more locally variable. Because this pattern appeared during both task engagement and aging, it may reflect increased local regulatory burden or reduced spatial coherence of hemodynamic propagation. However, the present data do not directly identify the underlying microvascular mechanisms. Therefore, fragmentation should be interpreted as an imaging marker of local hemodynamic irregularity rather than direct evidence of capillary- or arteriole-level regulation.

By contrast, changes in mean lag appear to capture a more global form of task-related hemodynamic tuning. In long-range white matter pathways, effective cognition may depend not on the magnitude of vascular reconfiguration alone, but on whether relative timing can be adjusted in a targeted and economical manner. Consistent with this interpretation, better cognitive performance was often associated with smaller task-rest lag differences, suggesting that high- performing individuals may require less extensive hemodynamic reconfiguration to meet task demands. Thus, ΔLag may provide a more behaviorally relevant index of system-level timing adjustment than local fragmentation alone.

This distinction also helps interpret the mediation results. Baseline WM markers were strongly related to age, but they did not independently explain cognitive performance after accounting for age. One possible explanation is that static baseline measures capture chronic physiological alteration, whereas cognition may depend more directly on the capacity for dynamic, task-evoked regulation. Under this interpretation, aging-related baseline fragmentation may indicate elevated resting-state load, but its behavioral consequences may only become evident when the system is challenged.

### A Baseline-Load Hypothesis for Aging-Related Hemodynamic Vulnerability

The distinction between system-level lag tuning and local spatial fragmentation provides a framework for interpreting the aging findings. Aging did not produce a simple global widening or narrowing of the white matter lag distribution. Instead, age-related effects were expressed locally within specific tracts, with increased spatial fragmentation and bidirectional shifts in relative mean lag. Importantly, many of these age-sensitive tracts overlapped with tracts that were modulated during working-memory task engagement in young adults. This anatomical concordance suggests that aging may alter the resting hemodynamic state of WM systems that are also recruited during cognitive demand.

These findings motivate a baseline-load hypothesis. Under this model, aging does not necessarily abolish dynamic vascular regulation. Rather, it may shift selected white matter systems toward a more irregular and higher-load resting configuration. If the baseline state is already closer to a task-engaged configuration, then subsequent cognitive demands may require less efficient, less targeted, or more extensive reconfiguration to achieve the same behavioral output. In this sense, aging may narrow the efficiency margin available for task-evoked hemodynamic tuning.

This interpretation should be viewed as a hypothesis rather than direct evidence of reduced vascular reserve. The present study demonstrates spatial convergence between task-sensitive white matter systems in young adults and age-sensitive resting-state alterations in older adults, but it does not directly measure task-evoked lag reconfiguration across the adult lifespan. Therefore, future studies combining resting-state and task-based fMRI in the same aging cohort will be necessary to test whether elevated baseline fragmentation predicts reduced task-evoked flexibility or poorer cognitive compensation.

### Limitations and Future Directions

Although Local Propagation Mapping reduces several artifacts associated with global-reference lag estimation, the present findings should be interpreted within several important limitations. First, fMRI provides only a macroscopic and indirect measure of cerebrovascular dynamics. Therefore, the observed spatial fragmentation of white matter lag cannot be assigned to specific cellular or microvascular mechanisms. Although increased fragmentation may reflect reduced spatial coherence or greater local regulatory burden, the present data do not directly identify the contributions of arterioles, capillaries, pericytes, glial signaling, or venous drainage. Future studies combining fMRI with higher-resolution vascular imaging, physiological challenges, or animal models will be necessary to clarify the biological sources of these lag patterns.

Second, the task and aging analyses were not performed as a single lifespan task-based experiment. Task-induced lag reconfiguration was measured in young adults, whereas aging-related alterations were assessed primarily from resting-state data across the adult lifespan. Thus, the observed overlap between task-sensitive and age-sensitive white matter systems provides indirect evidence for a baseline-load hypothesis, but it does not directly demonstrate that aging reduces task-evoked hemodynamic flexibility. Future studies should acquire both resting-state and task-based fMRI in the same aging cohort to test whether elevated baseline fragmentation predicts larger, less efficient, or less targeted task-evoked reconfiguration.

Third, the aging analysis was cross-sectional. As a result, the present study cannot determine whether baseline lag fragmentation precedes cognitive decline, emerges as a consequence of other age-related physiological changes, or reflects a compensatory adaptation. Longitudinal studies will be required to determine whether changes in white matter lag architecture track within-person cognitive trajectories over time. In addition, because LPM estimates relative timing within a centered lag field, future work should examine how these relative measures relate to absolute vascular transit time, perfusion, and cerebrovascular reactivity.

These limitations also define the next steps for translational research. Rather than treating static structural or resting-state markers as sufficient indicators of white matter health, future studies may benefit from incorporating dynamic stress tests that evaluate how white matter hemodynamics respond to cognitive demand. Such approaches could determine whether the combination of baseline lag organization and task-evoked reconfiguration provides a more sensitive marker of neurovascular vulnerability than either measure alone.

In conclusion, this study establishes a framework for mapping the spatiotemporal architecture of relative hemodynamic timing within human white matter. Using Local Propagation Mapping, we show that white matter lag patterns are spatially organized, reliable, and aligned with vascular anatomy. We further demonstrate that these patterns are modulated by cognitive state and that task-induced changes in macroscopic lag are associated with individual differences in cognitive performance. Across aging, resting-state white matter lag architecture shows tract-specific reorganization and increased spatial fragmentation, with partial overlap with task-sensitive systems observed in young adults. Together, these findings suggest that dynamic white matter hemodynamics may represent an important dimension of neurovascular regulation and motivate future lifespan studies testing whether elevated baseline load constrains task-evoked hemodynamic flexibility and cognitive resilience.

## METHODS

### Ethics Statement

The data involved in this research are publicly available and have been previously approved for use by the Washington University Institutional Review Board. All participants provided written informed consent to participate in this study. The authors did not collect any new data involving human participants.

### Dataset

We selected 687 entries from the Human Connectome Project Young Adult (HCP-Y)^38^ repository (comprising 335 males and 352 females, all between the ages of 22 and 35 years) and 688 subjects from HCP Aging (HCP-A)^39^ repository (308 males and 380 females, between ages of 36 and 100 years), adhering to criteria that included the completeness of a 3T scan and associated physiological data, along with adequate data quality. We have included only the relevant MRI modalities that were pertinent to our study from these databases. Those include resting state fMRI, working memory task fMRI, T1-weighted, and T2-weighted images. The sex of participants was recorded based on self-report in the HCP dataset. No sex- or gender-based analyses were conducted in this study.

For HCP-Y, the imaging protocols are detailed elsewhere^40^ and were performed with 3T Siemens Skyra scanners. fMRI scans were acquired using multiband gradient-echo EPI sequences. Each session consisted of two runs of scans with opposing phase encoding directions and lasted 14 minutes and 33 seconds, with parameters TR = 720 ms, TE = 33.1 ms, and an isotropic voxel resolution of 2 mm, totaling 1200 volumes for resting state and 405 volumes for working memory task. Concurrent recordings of physiological responses, such as respiration and heartbeat, were captured during fMRI scans. Additionally, T1-weighted images were obtained using a single-echo MPRAGE sequence with a TR of 2,400 ms, TE of 2.14 ms, and voxel dimensions of 0.7 mm isotropic. T2-weighted images were obtained using a 3D T2-SPACE sequence with a TR of 3200 ms, TE of 565 ms, and voxel dimensions of 0.7 mm isotropic.

For HCP-A, the imaging protocols are more thoroughly described elsewhere^41,42^. Briefly, scans were executed on Siemens 3T Prisma scanners with 32-channel head coils. The resting-state fMRI protocol involved four runs with opposing phase encoding directions, each 6 minutes and 41 seconds, with TR = 800 ms, TE = 37 ms, voxel dimension = 2 mm isotropic, and a total of 488 volumes for each run, while physiological parameters were also documented. T1-weighted images were obtained using a multi-echo MPRAGE sequence, with a TR of 2,500 ms, TEs of 1.8/3.6/5.4/7.2 ms, and voxel dimensions of 0.8 mm. T2-weighted images were obtained using a T2-SPACE sequence with a TR of 3200 ms, TE of 564 ms, and voxel dimensions of 0.8 mm isotropic.

### Preprocessing

Our approach was to employ ‘uncleaned’ images that underwent only the Minimal Preprocessing Pipelines^43^. Briefly, T1-weighted images were nonlinearly coregistered to MNI space using FNIRT^44^, with subsequent processing via the Freesurfer suite, resulting in voxel-wise BOLD time series^45^. Meanwhile, the T2-weighted images were aligned with the native T1-weighted images using 6 degrees of freedom (DOF) and subsequently registered to MNI space along with the T1-weighted images. The fMRI processing encompassed the removal of head movement artifacts, correction of distortions from susceptibility effects using FSL based on the two runs of data with opposite-phase encoding directions, and then nonlinear registration to MNI space. Further processing steps included regressing out confounding variables, including 12 head movement parameters and physiological fluctuations, modeled by the RETROICOR technique^46^. This preceded the application of linear trend corrections and temporal filtering using a band-pass filter covering the frequency range of 0.01 - 0.1 Hz.

### LPM Calculation

For every voxel within the brain mask, we extract the resting-state BOLD time series. We evaluate the pairwise temporal relationships strictly within a spatial neighborhood restricted to adjacent voxels. Because the temporal delay from a reference voxel to a target voxel is strictly the inverse of the delay in the opposite direction, we optimize computational efficiency by computing cross-correlations along thirteen independent spatial directions. For any given adjacent pair, we compute the discrete cross-correlation function across a temporal shift window spanning from negative to positive five repetition times.

The standard sampling rate of fMRI imaging inherently limits the temporal resolution of discrete correlation metrics. To achieve precise temporal mapping beyond the boundaries of the discrete sampling rate, we implement a three-point parabolic interpolation algorithm around the discrete peak of the correlation function^5,47^. Let *τ*_0_ represent the discrete temporal shift yielding the maximum correlation value r(*τ*_0_). Let r(*τ*_-1_) and r(*τ*_+1_) denote the correlation values at the immediately preceding and succeeding temporal shifts, respectively. The continuous peak shift *τ*_ij_ between voxel i and voxel j is mathematically calculated as

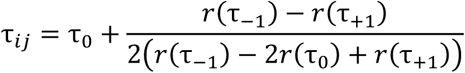

We establish a local directed edge from voxel i to voxel j exclusively if the discrete peak correlation avoids the temporal boundaries and exceeds a predefined reliability threshold of 0.15. The magnitude of this maximum correlation serves as the weight w_ij_ for the anatomical connection, mathematically reflecting the physiological reliability of the local lag estimation.

To synthesize these independent local delays into a coherent whole-brain topographical map, we conceptualize the entire brain architecture as a dense weighted graph. The fundamental analytical goal is to estimate a relative lag value T_i_ for every voxel such that the temporal difference between any adjacent pair optimally approximates the measured local delay *τ*_ij_. We frame this challenge as a least squares optimization problem. We aim to minimize the weighted sum of squared errors across all valid local edges, formulated as

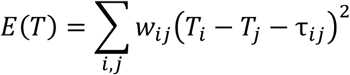

To minimize this global cost function, we compute the partial derivative with respect to the relative lag value of each individual voxel and equate the derivative to zero. This mathematical operation yields a linear equality for each node. The summation index j denotes the set of adjacent voxels within the immediate anatomical neighborhood of the reference voxel i according to the graph topology defined by the three-dimensional grid.

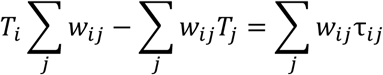

This massive set of interconnected linear equations elegantly transforms into a generalized graph Poisson equation

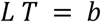

where matrix L represents the graph Laplacian defined as D - W. Matrix W contains all pairwise edge weights, and D represents a diagonal degree matrix where each diagonal element encapsulates the sum of weights connected to the corresponding voxel. The divergence vector b aggregates the weighted local delays for each voxel. We solve this extensive linear system using the preconditioned conjugate gradient algorithm to obtain the relative lag vector T. Because the solution is defined up to an additive constant, we subtract the spatial mean from the solution to center the temporal gradient. This step finalizes the generation of the topographical atlas for local hemodynamic propagation.

### Comparison with Alternative Lag Mapping Approaches and Physiological Validation

To evaluate the performance of Local Propagation Mapping against established neurovascular lag estimation techniques, we implemented two primary mapping approaches based on resting state fMRI. The first method utilizes global signal regression (GSR) lag mapping, which calculates the temporal shift of each voxel relative to the global mean signal of the brain. We extracted the global signal by averaging the time series across all intracranial voxels and performed Z-score normalization on both the global reference and individual voxel signals. We estimated the cross- correlation function within a shift range of five TR and applied a three-point parabolic interpolation to the discrete peak to achieve sub-TR precision. The second method employs global cross-correlation analysis, often referred to as the GCC approach, which determines the relative delay of a voxel by averaging the pairwise cross-correlations with every other voxel in the brain. This exhaustive pairwise calculation utilized the same parabolic interpolation refinement to ensure temporal accuracy.

The physiological validity of these lag estimations was assessed using an independent macroscopic venous partial volume atlas, known as the VENAT atlas^48^. This high-resolution reference was constructed from 7T quantitative susceptibility mapping (QSM) data acquired from young healthy volunteers. The atlas generation process involved repeated acquisitions to enhance the signal-to-noise ratio and utilized a multiscale vessel filter to segment the venous architecture. The resulting venous partial volume maps represent the density of macroscopic vessels and provide a biological ground truth for blood transit time distributions.

We performed several statistical evaluations to benchmark the reliability and specificity of the mapping techniques. To assess the spatial consistency between methods, we calculated the subject-wise spatial correlation (Pearson’s r) between the lag maps generated by each approach. To evaluate the internal stability of each algorithm, we performed a split-half reliability analysis. We randomly divided the subject cohort into two independent halves across 1,000 iterative permutations. For each permutation, we calculated the spatial intraclass correlation coefficient (ICC) between the group-averaged lag maps of the two halves. This iterative approach generates a distribution of reliability metrics to robustly quantify the estimation consistency. To investigate the alignment between hemodynamic delays and vascular anatomy, we compared the estimated lag values against the venous volume fractions. Because direct voxel-wise comparisons are heavily confounded by local signal noise and the non-normal distribution of signals across different tissue types, we parcellated the brain voxels into contiguous spatial bins. We sorted all valid voxels based on their venous volume fractions and divided them into fifty equal-sized bins. We then computed the average hemodynamic delay and the average venous volume for each bin. To characterize the relationship between regional venous density and the estimated delays, we applied a simple linear regression model to these bin-averaged data points. We extracted the coefficient of determination, denoted as the R^2^ statistic, to quantify the proportion of variance in the hemodynamic lag that is linearly explained by the underlying macroscopic venous density. Furthermore, we quantified the separation sensitivity of each method by examining the extremes of the venous density distribution. For every individual subject, we extracted the mean estimated delay within a spatial mask representing regions of extremely high venous density, defined as the upper quartile (top 25%) of the venous volume distribution. We similarly extracted the mean delay within regions of extremely low venous density, defined as the lower quartile (bottom 25%). To determine the capacity of each method to differentiate the physiological signatures of distinct vascular environments, we computed Cohen’s d effect size. We calculated this metric by taking the mean difference in delays between the upper and lower quartiles and dividing it by the pooled standard deviation of the delays across the respective regions. This standardized effect size provides a robust statistical measure of physiological specificity.

### Spatial Profiling of Hemodynamic Lag and Metric Quantification

To spatially profile the hemodynamic propagation within specific neural pathways, we utilized a population-averaged atlas of the macroscopic human structural connectome ^49^. This high angular resolution brain atlas was constructed using diffusion magnetic resonance imaging data from one thousand and sixty-five subjects. We employed DSI Studio software to project the estimated voxel-wise hemodynamic delay maps onto the trajectories of seventy major WM bundles. These bundles encompass the primary association commissural and projection pathways of the human brain. For each specific anatomical bundle, the along-tract analysis extracted the hemodynamic lag signals at one hundred equidistant nodes spanning from the origin to the termination of the tract. This procedure effectively maps the three-dimensional voxel-wise delays into a standardized one-dimensional spatial gradient along the axonal architecture.

To comprehensively quantify the spatiotemporal architecture of these hemodynamic gradients and their modulations, we computed three distinct metrics. First, to assess the global temporal dispersion of blood flow across the entire WM, we extracted the voxel-wise lag distribution and calculated the FWHM. A wider distribution indicates an expansion of the macroscopic dynamic range of the neurovascular system. Second, at the individual tract level, we calculated the macroscopic delay by averaging the transit time values across all one hundred nodes within the tract. This macroscopic delay metric reflects the relative overall temporal shift or the functional throughput of blood flow within the specific neural pathway. Third, to capture the microscopic spatial complexity of the propagation pattern, we performed a spatial spectral analysis on the demeaned along-tract lag profile. Because the spatial power spectrum of these hemodynamic gradients typically follows a power law distribution, the relationship exhibits a near-linear trend when transformed into a log-log coordinate system. We therefore estimated the spatial spectral slope by applying a linear regression model to the logarithm of the spatial frequency and the logarithm of the power. We quantified the spatial fragmentation through this slope, where a shallower spectral slope indicates an increased presence of high-frequency spatial fluctuations representing a more fragmented local microvascular regulation.

### Assessment of Task-Induced Hemodynamic Modulation and Cognitive Associations

To quantify the task-induced global modulation of the WM dynamic range, we compared the FWHM of the whole-brain voxel-wise lag distributions between the resting state and the task state. Furthermore, we evaluated the corresponding modulations of the macroscopic delay and the spatial fragmentation at the level of individual anatomical bundles. For these comparisons, we calculated the within-subject difference between the task and resting states for each metric. We applied a linear regression framework to evaluate the significance of these state transitions. To isolate the genuine physiological modulation from potential scanning artifacts, we included the state difference in in-scanner head motion as a continuous covariate in all models. The statistical significance of the state transition was determined by the intercept of the regression model which represents the group-level task effect after adjusting for motion differences. We applied a false discovery rate procedure to correct the resulting probability values for multiple comparisons across the seventy anatomical tracts.

To investigate the behavioral relevance of task-induced hemodynamic reconfiguration, we examined the statistical associations between task-induced modulations of WM hemodynamics and individual cognitive performance. Specifically, we evaluated whether dynamic shifts in macroscopic delay and spatial fragmentation could independently relate to cognitive abilities across multiple domains. We extracted six distinct behavioral metrics, including the fluid intelligence composite score, the progressive matrices accuracy, the dimensional change card sort performance, the processing speed, the list sorting working memory score, and the accuracy of the two-back working memory task. We constructed multiple linear regression models in which each cognitive score served as the dependent variable and the task-induced modulation of either macroscopic delay or spatial fragmentation served as the primary predictor. To rigorously isolate the specific behavioral associations, we incorporated chronological age, biological sex, the corresponding baseline resting-state hemodynamic metric, and the state difference in head motion as covariates within each regression model. We determined the significance of the behavioral association based on the independent statistical contribution of the task-induced modulation term, and we adjusted the final probability values using false discovery rate correction across the investigated tracts within each cognitive domain.

### Statistical Modeling of Aging and Mediation Pathways

To investigate the influence of chronological age on WM hemodynamic lags, we utilized the HCP-A dataset. We excluded individuals with a recorded age exceeding ninety-nine years to ensure the stability of the statistical estimations. We constructed multiple linear regression models to evaluate the age-related changes in the macroscopic delay and the spatial fragmentation within each of the seventy anatomical tracts. In these models, the specific hemodynamic metric served as the dependent variable while chronological age was the primary predictor of interest. To isolate the genuine effect of aging, we incorporated biological sex and the mean relative head motion as nuisance covariates. We determined the statistical significance of the age effect based on the t statistic of the age term, and we applied the false discovery rate procedure to adjust for multiple comparisons across the entire set of fiber bundles.

To assess the relationship between hemodynamic propagation and structural tissue properties, we quantified the regional myelin content through the ratio of T1-weighted and T2-weighted images. We applied a voxel-wise division of the intensity restored T1-weighted image by the T2-weighted image and implemented an intensity capping procedure to minimize the impact of noise artifacts in the WM. Using the DSI Studio framework, we projected these myelin maps onto the same seventy anatomical tracts and extracted the average myelin content across one hundred nodes per tract to ensure spatial correspondence with the hemodynamic metrics.

We further employed a formal mediation analysis framework to explore the sequential relationships between chronological age, the neuroimaging markers, and cognitive performance. This framework evaluated three distinct pathways within a single model. Path (a) quantified the effect of age on the brain markers, including macroscopic delay, lag fragmentation, and myelin content. Path (b) assessed the independent predictive value of each brain marker for cognitive scores after statistically regressing out the variance of chronological age. We utilized the Sobel test to calculate the overall mediation strength and to determine if the biological markers significantly mediated the impact of aging on cognitive functions. All paths in the mediation models were adjusted for sex and head motion to ensure the robustness of the identified associations.

## Data Availability

The HCP-Y data used in this study are available in the HCP database https://www.humanconnectome.org/

The HCP-A data used in this study are available in the NIMH Data Archive https://nda.nih.gov

## Code Availability

The custom scripts and the computational algorithms developed for the neuroimaging analysis in this study are freely available in the public GitHub repository at: https://github.com/geyerou/White-Matter-Lag

Detailed instructions for reproducing the key results are provided in the repository’s README file.

Other software and toolboxes that are required for running the code include:

Nilearn: https://nilearn.github.io/

GIFTI: https://www.nitrc.org/projects/gifti/

DSI Studio: https://dsi-studio.labsolver.org/

SPM12: https://www.fil.ion.ucl.ac.uk/spm/software/spm12/

NIFTI Toolbox: https://github.com/mcnablab/NIFTI_toolbox

## ACKNOWLEDGMENTS

This work was supported by the National Institutes of Health (NIH) grants R01 NS113832 (JG) and R01 NS129855 (ZD).

Data of young adults were provided by the Human Connectome Project, WU-Minn Consortium (Principal Investigators: David Van Essen and Kamil Ugurbil; 1U54MH091657), funded by the 16 NIH Institutes and Centers that support the NIH Blueprint for Neuroscience Research; and by the McDonnell Center for Systems Neuroscience at Washington University.

HCP-Aging data reported in this publication were supported by the National Institute on Aging of the National Institutes of Health under Award Number U01AG052564 and by funds provided by the McDonnell Center for Systems Neuroscience at Washington University in St. Louis. The HCP-Aging 2.0 Release data used in this report came from DOI: 10.15154/1520707.

## AUTHOR CONTRIBUTIONS

M.L.: Writing, Visualization, Validation, Software, Methodology, Investigation, Conceptualization. Z.D.: Supervision and Funding acquisition. J.C.G: Supervision and Funding acquisition.

