## Supplemental Figures for "Relative Hemodynamic Timing in Human White Matter"

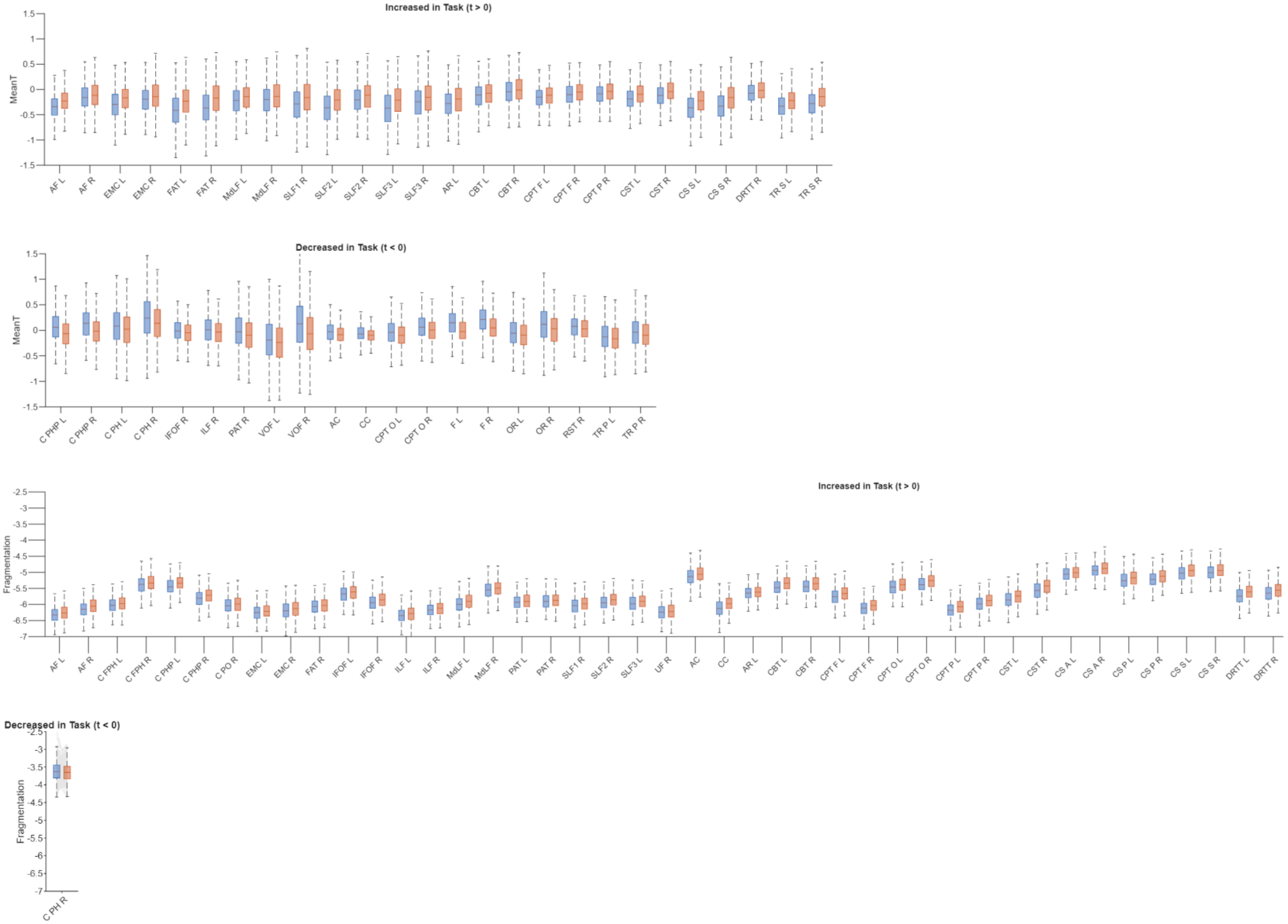


**Figure S1. Exhaustive profiling of task-induced hemodynamic modulations across all significant white matter tracts.** Extended results complementing Figure 3 in the main text. Group-level comparisons of hemodynamic properties between resting-state (blue boxes) and task-state (orange boxes) are shown for the complete set of white matter tracts that survived rigorous statistical thresholding (FDR-corrected $P < 0.05$, adjusting for state-specific head motion). The panels display comprehensive distributions for two metrics: macroscopic delay (Mean T), separated into tracts with significant task-induced increases (top panel, $t > 0$) and decreases (second panel, $t < 0$); and spatial fragmentation (spectral slope $\beta$), separated into significant task-driven increases (third panel) and decreases (bottom panel). Note that while widespread increases in spatial fragmentation are observed across the white matter architecture, only a single tract (C PH R) exhibits a significant decrease. Anatomical abbreviations on the x-axis correspond to the standard 70-tract DSI Studio atlas.

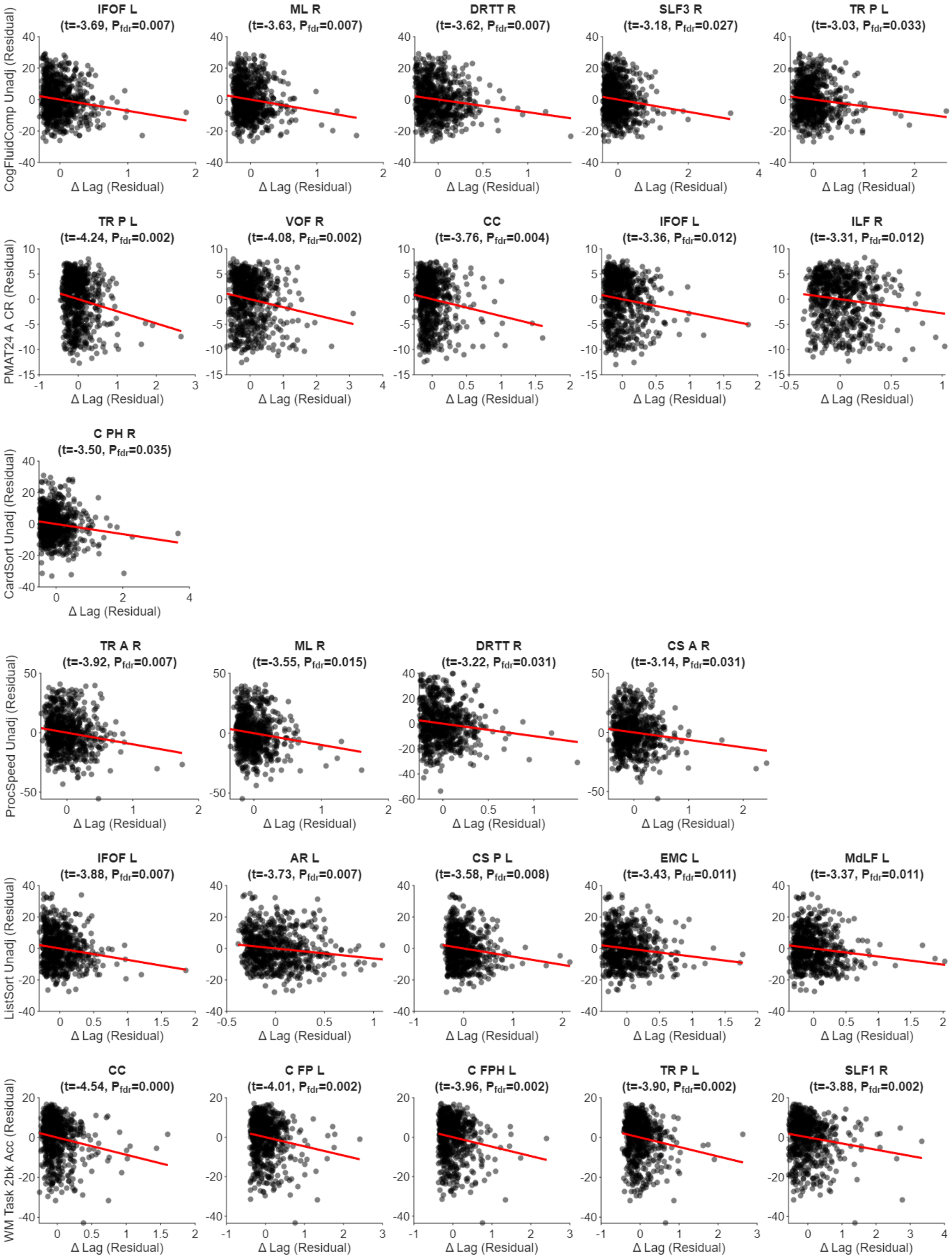


**Figure S2. Partial regression plots showing the association between hemodynamic delay modulation (Delta Lag) and cognitive performance.** Each row presents scatter plots for a specific cognitive domain (from top to bottom: Fluid Intelligence Composite, Penn Matrix Test, Card Sorting, Processing Speed, Working Memory/List Sorting, and 2-back Task Accuracy). For each domain, the top 5 most significantly associated white matter tracts (ranked by T-statistic) are shown. Both the x-axis (Delta Lag) and y-axis (Cognitive Score) represent residuals from linear models that partial out the effects of age, gender, baseline lag, and head motion difference (Delta Motion), thus illustrating the unique relationship between hemodynamic flexibility and behavior. Each point represents an individual subject. The red line indicates the linear regression fit. Tract names, T-statistics, and FDR-corrected P-values ($P_{FDR}$) are provided above each plot.

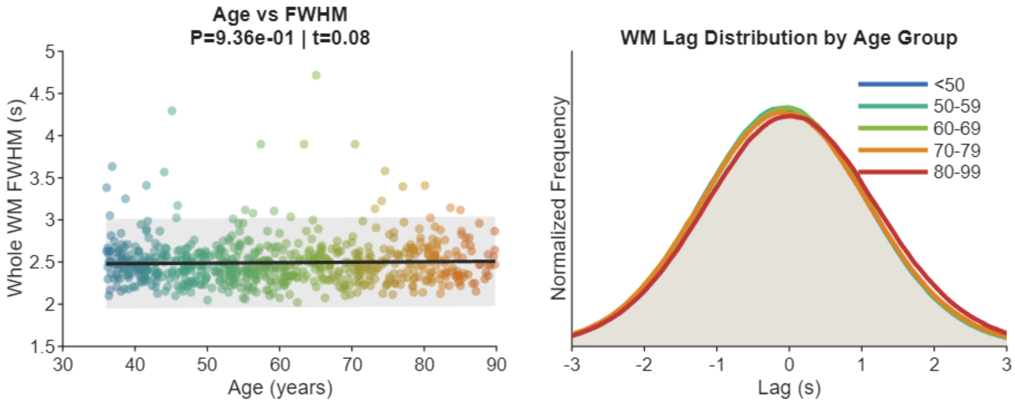


**Figure S3. Stability of the macroscopic hemodynamic delay gradient over aging.** (Left) Age versus Whole White Matter FWHM. Scatter plot illustrating the relationship between chronological age and the Full Width at Half Maximum (FWHM) of the whole-brain white matter lag distribution. Each dot represents an individual subject, color-coded by their respective age cohort. The solid black line indicates the linear regression fit, with the 95% confidence interval shaded in light grey. No significant age-related change was observed in the overall dynamic range of the lag distribution (P = 0.936, t = 0.08), after controlling for sex and head motion. (Right) Hemodynamic delay distributions across age groups. Normalized voxel-wise frequency histograms of the white matter lag distribution, averaged within five age cohorts (from <50 to 80-99 years). The near-perfect overlap of these distributions demonstrates that the global spatial dispersion and temporal architecture of the macroscopic hemodynamic gradient are highly conserved and robust against chronological aging.
