## Supplemental Tables for "Relative Hemodynamic Timing in Human White Matter"

Table S1. Abbreviations of the tract names

| **Association** | | **Projection** | |
| --- | --- | --- | --- |
| AF_L | Arcuate_Fasciculus_L | AR_L | Acoustic_Radiation_L |
| AF_R | Arcuate_Fasciculus_R | AR_R | Acoustic_Radiation_R |
| C_FPH_L | Cingulum_Frontal_Parahippocampal_L | CBT_L | Corticobulbar_Tract_L |
| C_FPH_R | Cingulum_Frontal_Parahippocampal_R | CBT_R | Corticobulbar_Tract_R |
| C_FP_L | Cingulum_Frontal_Parietal_L | CPT_F_L | Corticopontine_Tract_Frontal_L |
| C_FP_R | Cingulum_Frontal_Parietal_L | CPT_F_R | Corticopontine_Tract_Frontal_R |
| C_PHP_L | Cingulum_Parahippocampal_Parietal_L | CPT_P_L | Corticopontine_Tract_Parietal_L |
| C_PHP_R | Cingulum_Parahippocampal_Parietal_R | CPT_P_R | Corticopontine_Tract_Parietal_R |
| C_PH_L | Cingulum_Parahippocampal_L | CPT_O_L | Corticopontine_Tract_Occipital_L |
| C_PH_R | Cingulum_Parahippocampal_R | CPT_O_R | Corticopontine_Tract_Occipital_R |
| C_PO_L | Cingulum_Parolfactory_L | CS_A_L | Corticostriatal_Tract_Anterior_L |
| C_PO_R | Cingulum_Parolfactory_R | CS_A_R | Corticostriatal_Tract_Anterior_R |
| EMC_L | Extreme_Capsule_L | CS_S_L | Corticostriatal_Tract_Superior_L |
| EMC_R | Extreme_Capsule_R | CS_S_R | Corticostriatal_Tract_Superior_R |
| FAT_L | Frontal_Aslant_Tract_L | CS_P_L | Corticostriatal_Tract_Posterior_L |
| FAT_R | Frontal_Aslant_Tract_R | CS_P_R | Corticostriatal_Tract_Posterior_R |
| IFOF_L | Inferior_Fronto_Occipital_Fasciculus_L | CST_L | Cortico_Spinal_Tract_L |
| IFOF_R | Inferior_Fronto_Occipital_Fasciculus_R | CST_R | Cortico_Spinal_Tract_R |
| ILF_L | Inferior_Longitudinal_Fasciculus_L | DRTT_L | Dentatorubrothalamic_Tract_L |
| ILF_R | Inferior_Longitudinal_Fasciculus_R | DRTT_R | Dentatorubrothalamic_Tract_R |
| MdLF_L | Middle_Longitudinal_Fasciculus_L | F_L | Fornix_L |
| MdLF_R | Middle_Longitudinal_Fasciculus_R | F_R | Fornix_R |
| PAT_L | Parietal_Aslant_Tract_L | ML_L | Medial_Lemniscus_L |
| PAT_R | Parietal_Aslant_Tract_R | ML_R | Medial_Lemniscus_R |
| SLF1_L | Superior_Longitudinal_Fasciculus1_L | OR_L | Optic_Radiation_L |
| SLF1_R | Superior_Longitudinal_Fasciculus1_R | OR_R | Optic_Radiation_R |
| SLF2_L | Superior_Longitudinal_Fasciculus2_L | RST_L | Reticulospinal_Tract_L |
| SLF2_R | Superior_Longitudinal_Fasciculus2_R | RST_R | Reticulospinal_Tract_R |
| SLF3_L | Superior_Longitudinal_Fasciculus3_L | TR_A_L | Thalamic_Radiation_Anterior_L |
| SLF3_R | Superior_Longitudinal_Fasciculus3_R | TR_A_R | Thalamic_Radiation_Anterior_R |
| UF_L | Uncinate_Fasciculus_L | TR_P_L | Thalamic_Radiation_Posterior_L |
| UF_R | Uncinate_Fasciculus_R | TR_P_R | Thalamic_Radiation_Posterior_R |
| VOF_L | Vertical_Occipital_Fasciculus_L | TR_S_L | Thalamic_Radiation_Superior_L |
| VOF_R | Vertical_Occipital_Fasciculus_R | TR_S_R | Thalamic_Radiation_Superior_L |
| **Commissural** | |  |  |
| AC | Anterior_Commissure |  |  |
| CC | Corpus_Callosum |  |  |
